# Modeling and quantifying frequency-dependent fitness in microbial populations with cross-feeding interactions

**DOI:** 10.1101/012807

**Authors:** Noah Ribeck, Richard E. Lenski

## Abstract

Coexistence of multiple populations by frequency-dependent selection is common in nature, and it often arises even in well-mixed experiments with microbes. If ecology is to be incorporated into models of population genetics, then it is important to represent accurately the functional form of frequency-dependent interactions. However, measuring this functional form is problematic for traditional fitness assays, which assume a constant fitness difference between competitors over the course of an assay. Here, we present a theoretical framework for measuring the functional form of frequency-dependent fitness by accounting for changes in abundance and relative fitness during a competition assay. Using two examples of ecological coexistence that arose in a long-term evolution experiment with *Escherichia coli*, we illustrate accurate quantification of the functional form of frequency-dependent relative fitness. Using a Monod-type model of growth dynamics, we show that two ecotypes in a typical cross-feeding interaction—such as when one bacterial population uses a byproduct generated by another—yields relative fitness that is linear with relative frequency.

## Introduction

The coexistence of diverse organisms is a key issue in both ecology and evolutionary biology. While much of the diversity of life is undoubtedly maintained by geographic isolation and niche specialization, stable coexistence can also occur by more subtle interactions between distinct ecotypes within a local population. In some cases, this ecotypic diversity persists even in a well-mixed, constant environment with only a single exogenously supplied resource, in apparent violation of the competitive exclusion principle (Gause 1934; Hardin 1960). Such polymorphisms are maintained by negative frequency-dependent selection, where each type persists at some intermediate frequency owing to a selective advantage when rare (Antonovics and Kareiva 1988).

Polymorphisms sustained by frequency dependence have been observed and quantified in many systems, including genes associated with the immune response (Borghans et al. 2004), chromosomal inversions in *Drosophila* (Wright and Dobzhansky 1946), inbred lines of *C. elegans* (Chelo et al. 2013), and color polymorphisms in lizards (Sinervo and Lively 1996). Laboratory experiments with well-mixed populations of microbes have frequently generated and sustained stable polymorphisms that have been shown to involve balancing selection resulting from frequency dependence (Levin 1972; Helling et al. 1987; Rosenzweig et al. 1994; Turner et al. 1996; Elena and Lenski 1997; Treves et al. 1998; Rainey et al. 2000; Rozen and Lenski 2000; Blount et al. 2008). These interactions play an important role in the dynamics of microbial populations because ecotypic polymorphisms are robust and widespread (Pfeiffer and Bonhoeffer 2004; Estrela and Gudelj 2010), and they drive diversification even in sympatry (Dieckmann and Doebeli 1999; Doebeli and Dieckmann 2000; Friesen et al. 2004; Rozen et al. 2009; Herron and Doebeli 2013).

Despite the importance of ecological interactions in adaptation, many models of evolution ignore them. For example, Fisher’s fundamental theorem of natural selection (Fisher 1930), the Price equation (Price 1970), and the theory of adaptation in large asexual populations (Desai and Fisher 2007) all ignore frequency-dependent selection. To understand adaptation more generally, one must interpret ecological interactions in the context of fitness, which is the ultimate metric that governs the contribution of different genotypes to future generations. Among other issues, one must know the functional form of frequency-dependent fitness in populations with coexisting lineages to interpret and understand their evolutionary dynamics.

To that end, we aim to characterize quantitatively how the relative fitness of coexisting ecotypes depends on their relative frequency. However, traditional fitness assays infer relative fitness from changes in the relative frequency of two competitors over time, which assumes a constant fitness differential between them (Lenski 1988; Lenski et al. 1991). To remedy this problem, we describe a procedure for measuring frequency-dependent selection that accounts for the dynamically changing abundances and hence relative fitness over the course of a fitness assay. We demonstrate this procedure using two examples of ecotypic divergence and coexistence that arose in a long-term evolution experiment with *E. coli* (Lenski et al. 1991; Rozen and Lenski 2000; Le Gac et al. 2012).

To put these measurements into context, we then use a model of growth dynamics to examine cross-feeding interactions, which are a common source of negative frequency-dependent selection in evolving microbial populations (Pfeiffer and Bonhoeffer 2004; Estrela and Gudelj 2010). In this case, one ecotype has an inferior ability to utilize the exogenously supplied resource that limits the overall density of the population, but it is able to persist by more efficiently using a metabolic byproduct that one or both ecotypes produce (Doebeli 2002). For this scenario, we model the Monod dynamics of resource use and population growth, and we calculate the resulting frequency-dependent form of the relative fitness of the two types. We find that the relative fitness for two ecotypes in a cross-feeding interaction is generally linear with respect to relative frequency.

## Methods

### Competition assays

Competitions were performed in the same DM25 medium used in the *E. coli* long-term evolution experiment (LTEE) using the procedures described in detail elsewhere (Lenski et al. 1991). Multiple growth cycles were achieved by diluting cultures 100-fold (dilution factor *F* = 100) into fresh medium each day. To avoid large sampling error resulting from small counts, competitions with extreme initial ratios (*p*_0_ ≤ 0.1 and *p*_0_ ≥ 0.9) were plated three times, those with intermediate ratios (0.1 < *p*_0_ < 0.5 and 0.5 < *p*_0_ < 0.9) were plated twice, and those started with equal ratios (*p*_0_ = 0.5) were plated once.

### S and L competitors

Two persistent ecotypes initially diverged in the LTEE Ara–2 population after ∼6,000 generations. The two types are denoted S and L (small and large) on the basis of their colony sizes on agar plates: S colonies are distinctively smaller and barely visible after 24 h (Rozen and Lenski 2000). Single clones from the S and L ecotypes were obtained from the archived sample of Ara–2 that was frozen at 6,500 generations. The clones were isolated by overnight growth in LB medium, spreading cells onto an agar plate, and incubating them for 48 h at 37°C. At 6,500 generations, the S ecotype was unable to utilize maltose, but most L cells retained the ability to grow on maltose; a Mal^+^ L clone was chosen for this study. To measure relative frequencies in competitions, the S and L colonies were distinguished on the basis of their red and white coloration, respectively, after incubation on a tetrazolium maltose (TM) agar plate for 48 h at 37°C. We performed six replicate competitions between the S and L clones at each of seven different initial relative frequencies. As an estimate of the sampling error associated with the plating procedures, the standard deviation of the measured relative frequency among multiply plated competitions is *σ*_*p*_ = 0.023. Confidence intervals on fit parameters were obtained by performing 20 bootstrap-resampled curve fits.

### *nuoM*^+^ and *nuoM*^−^ competitors

We used 22 clones previously chosen at random from the Ara–1 population sample at 10,000 generations (Elena and Lenski 1997). Previous work indicated that a particular mutation at the *nuoM* locus existed at an intermediate frequency in this sample (Barrick and Lenski 2009), and Sanger sequencing was performed with each clone to determine whether it carried the ancestral (*nuoM*^+^) or mutant (*nuoM*^−^) allele. The 8 clones identified as *nuoM*^−^ were grown overnight in LB medium, and equal volumes were combined to make an admixture. From each of the 14 clones identified as *nuoM*^+^, we used a spontaneous Ara^+^ mutant that can grow on arabinose (Elena and Lenski 1997); these 14 Ara^+^ derivatives were grown and mixed together in the same way. Ara^+^ mutants have been shown to be selectively neutral in the medium of the LTEE (Lenski et al. 1991; Elena and Lenski 1997), although that fact is not essential for this study. To measure initial and final relative frequencies, colonies were distinguished on the basis of the arabinose-utilization marker after incubation on tetrazolium arabinose (TA) agar plates for 24 h at 37°C. Three replicate competitions were done at each of seven initial relative frequencies. As an estimate of the sampling error associated with the plating procedures, the standard deviation of the measured relative frequency among multiply plated competitions is *σ*_*p*_ = 0.017. Confidence intervals on fit parameters were obtained by performing 20 bootstrap-resampled curve fits.

### Model of cross-feeding interaction

Using Wolfram Mathematica (version 8.0), we obtained numerical solutions of a system of Monod equations that describe the cross-feeding interaction in a serial-transfer culture regime. The final population sizes of each ecotype were determined at time points after both resources were depleted prior to the next transfer (it is assumed that there is no mortality in either ecotype).

## Results

### Quantifying frequency dependence

In experiments with microbes, relative fitness is typically measured by the change in relative frequencies of two marked strains placed in competition (Lenski 1988; Lenski et al. 1991). However, frequency-dependent interactions confound this measurement, because relative fitness changes over the course of the assay as the competitors’ frequencies change. To address this problem, we present a theoretical framework for measuring the functional form of the frequency-dependent fitness of two competing types within an asexual population.

For two coexisting ecotypes, the relative fitness must be characterized by negative frequency dependence, where each type has a competitive advantage when rare. The fitness must therefore decrease monotonically with relative frequency, which can be described most simply by a decreasing line, which we parameterize with intercept 1 + *s* and slope −*ms*. Then, the frequency-dependent relative fitness *w*(*p*) and population mean fitness 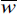 are given by:

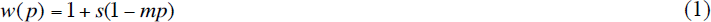

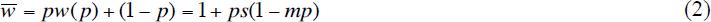

The equilibrium frequency is *p*_eq_ = 1/*m* (such that *w*(*p*) = 1), while the magnitude of the slope *ms* describes the strength of frequency dependence. The ecotype frequency dynamics can then be described by:

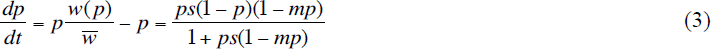

If the strength of selection is not extremely strong (*s* ≪ 1 and *s*(*m* − 1) ≪ 1), this can be approximated as:

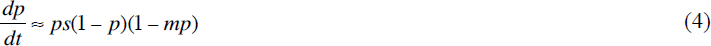

Separation of variables and integration then provides a formula that relates parameters *m* and *s* to the initial and final relative frequencies *p*_0_ and *p*_*f*_ obtained from competition assays:

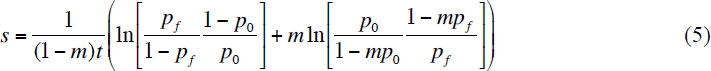

When frequency dependence is absent (*m* = 0), the right side reduces to the traditional expression for log-fitness (Chevin 2011). Competition assays may be performed over multiple serial transfers, so the competition time in generations is *t* = *k* log_2_(*F*), where *k* is the number of transfers and *F* is the dilution factor for each transfer. To estimate the frequency-dependent fitness, we seek values for the unknown parameters *m* and *s* that minimize the mean-squared difference between measured *p*_*f*_ and the theoretical value of *p*_*f*_ that satisfies Eq. 5. However, since Eq. 5 can not be solved explicitly for *p*_*f*_, numerical solutions must be used for this curve-fitting procedure. While a unique solution of the two parameters *m* and *s* requires at least two measurements at different sets of *p*_0_ and *p*_*f*_, the ideal experimental design consists of many competitions across a wide range of initial frequencies, with final frequencies measured at multiple time points (optimal experimental design is discussed further below).

To illustrate this method, we use fitness assays to measure the frequency-dependent fitness of pairs of clones from two independent LTEE populations that have evolved genotypes that are able to stably coexist. In one case, using clones isolated at 10,000 generations from population Ara–1, we found that individuals with a particular mutation in the *nuoM* gene (*nuoM*^−^) (Barrick and Lenski 2009) exhibit frequency-dependent selection in competition with individuals with the wild-type allele (*nuoM*^+^). The *nuoM*^−^ ecotype persisted at an intermediate frequency for ∼7,500 generations (Barrick and Lenski 2009; Barrick et al. 2009). To measure the fitness of the *nuoM*^−^ type relative to the *nuoM*^+^ type as a function of the *nuoM*^−^ frequency *p*, we completed a series of competitions (*k* = 6 daily transfers, *F* = 100 fold daily growth) with varying initial frequencies (Fig. 1). Applying these data to Eq. 5 produces estimates of *s* = 0.004 ± 0.003 and *m* = 9 ± 4 (95% CI). These estimates in turn indicate an equilibrium frequency of the *nuoM*^−^ ecotype *p*_eq_ = 0.11 ± 0.06 (95% CI), which accords well with the ∼16% frequency of *nuoM*^−^ at 10,000 generations that was estimated by metagenomic sequencing (Barrick and Lenski 2009) Note that these competing populations are admixtures of clones isolated from the 10,000 generation mixed population (see Methods), so the precise relationship between the two ecotypes in the evolved population may be slightly different.

**Figure 1.**
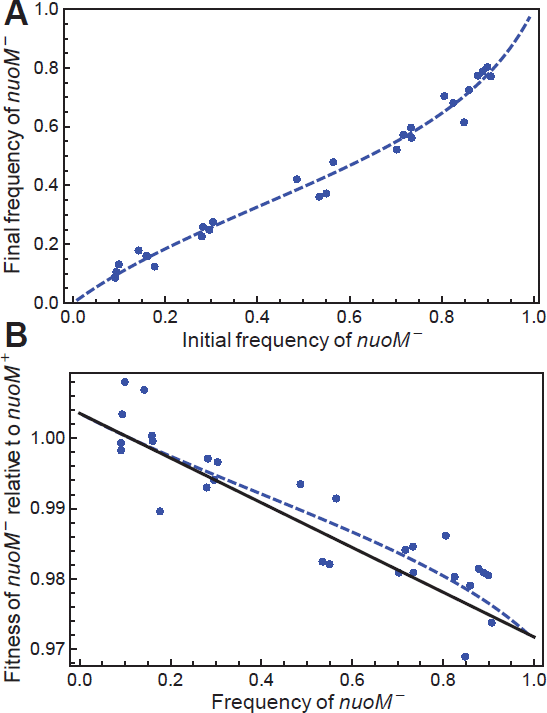
Experimental case of frequency dependence involving *nuoM* mutants. (A) Results from competition assays between mutant (*nuoM*^−^) and wild-type (*nuoM*^+^) clones. Blue points indicate relative frequency after 6-day competition as a function of initial frequency. The blue dashed curve indicates the best-fit model of linear frequency dependence (see Eq. 5). (B) Frequency-dependent fitness in the *nuoM* system. The black curve indicates the best-fit model of linear frequency dependence *w*(*p*) calculated in panel A. The blue points indicate mean fitness over the course of the 6-day competition, without adjusting for the dynamically changing relative fitness. The blue dashed curve shows the 6-day mean fitness prediction generated by *w*(*p*).

In the second case, two lineages denoted S and L initially diverged in one of the LTEE populations after ∼6,000 generations (Rozen and Lenski 2000) and have continued to coexist at least through 50,000 generations by a cross-feeding interaction (Le Gac et al. 2012; Plucain et al. 2014). To measure the fitness of S relative to L as a function of frequency *p*, we competed clones of each type isolated from 6,500 generations with varying *p*_0_, and measured *p*_*f*_ after 6 and 12 days of serial dilution and 100-fold growth. In this case, however, the set of competition assays is not consistent with a single linear parameterization of frequency dependence (Fig. S1). Instead, an alternate descriptive model for *w*(*p*) must be considered. We find that a quadratic representation *w*(*p*) = 1 + *s*(1 − *mp* + *qp*^2^) accurately generates the 6-day and 12-day mean fitness measurements for each *p*_0_, with best-fit parameters *s* = 0.026 ± 0.007, *m* = 6.5 ± 0.5, and *q* = 3.3 ± 0.9 (95% CI) (see Fig. 2; ΔAIC = 15.3, odds ratio = 2130). The resulting fit gives an equilibrium frequency of the S ecotype *p*_eq_ = 0.17 ± 0.02, which is consistent with the frequency of S estimated at 6,500 generations based on colony morphology (Rozen and Lenski 2000).

**Figure 2.**
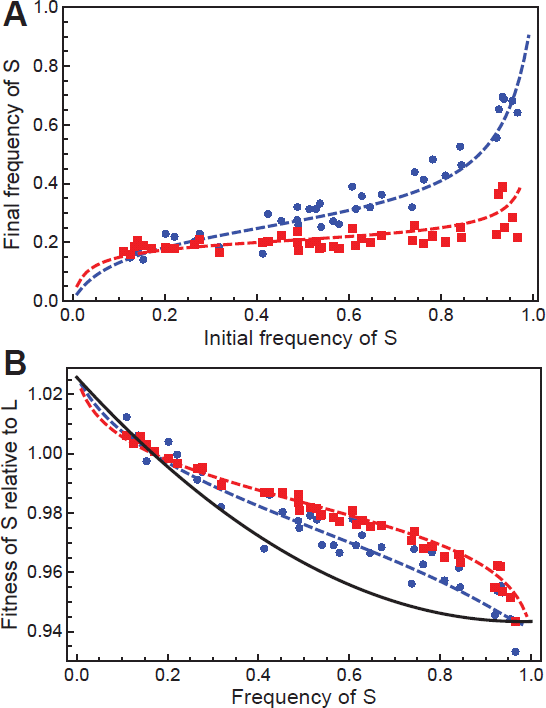
Experimental case of frequency dependence involving S and L ecotypes. (A) Results from competition assays between S and L clones. Blue circles and red squares indicate relative frequency after 6-day and 12-day competition, respectively, as a function of initial frequency. The blue dashed and red dashed curves indicate the best fit of the combined 6-day and 12-day data to a model of quadratic frequency dependence. (B) Frequency-dependent fitness of the S and L ecotypes. The black curve indicates the best-fit model of quadratic frequency-dependent fitness *w*(*p*) calculated in panel A. The blue and red points indicate mean fitness over the course of 6-day and 12-day competition, respectively, without adjusting for the dynamically changing relative fitness. The blue dashed and red dashed curves show the respective 6-day and 12-day mean fitness predictions generated by *w*(*p*).

In general, this curve-fitting procedure can be done for any parameterization of *w*(*p*), and the resulting model can be solved by fitting competition data to a numerical calculation of *p*_*f*_ for each *p*_0_. This procedure requires as least as many competitions as there are free parameters in the representation of *w*(*p*).

### Model of frequency dependence with cross-feeding interaction

To this point, we have focused on a procedure for measuring frequency-dependent fitness in microbial populations, assuming a linear parameterization. To evaluate the appropriateness of this assumption, we must identify the expected functional form of frequency-dependence for typical ecological interactions. For evolving microbial populations, a common source of negative frequency-dependent selection is cross-feeding (Pfeiffer and Bonhoeffer 2004; Estrela and Gudelj 2010). In this type of interaction, one ecotype has a superior ability to utilize a primary resource, while the other has a superior ability to utilize a metabolic byproduct that one or both ecotypes produce. To calculate the expected frequency dependence from such an ecological interaction, we use a model of resource use and population growth based on a Monod model (Doebeli 2002). Many evolution experiments (including those analyzed here) are performed in batch culture with periodic (e.g., daily) transfers to fresh media, so we model the growth of two ecotypes as they compete for a single exogenously supplied resource that is depleted between transfers. This model also includes a second, endogenously generated resource that is secreted, re-assimilated, and used for further growth by one or both types. By modeling the dynamics of competition across a range of initial relative frequencies, we can find final relative frequencies after each growth cycle and compute the functional form of the frequency-dependent fitness between the two ecotypes.

The key to coexistence under this scenario is the production and secretion of the secondary resource (e.g., acetate). This byproduct effectively provides a second ecological niche, where a genotype with an inferior ability to grow on the primary resource (e.g., glucose) may be able to persist if it is a better competitor for the secondary resource. Consider first a simple, hypothetical case, where one ecotype grows exclusively on a finite supply of glucose, and the other grows exclusively on acetate byproduct. In this case, provided enough time, each type will simply grow to the carrying capacity of its respective energy source, regardless of the initial ratio of the two subpopulations. This final relative frequency represents the ecological equilibrium of the mixed population. If either ecotype is initially rare (i.e., starts at below the equilibrium ratio), then it will increase in frequency and thus appear to have a fitness advantage until the equilibrium ratio is achieved. More generally, there will be negative frequency-dependent fitness between two ecotypes provided that each type exhibits superior growth on one of the resources. A stable equilibrium ratio exists if the abilities of the two types are sufficiently balanced, although it may take multiple cycles of dilution and growth for the population to achieve that equilibrium.

To calculate the frequency-dependent fitness of two ecotypes in a cross-feeding interaction, we model their population growth using Monod dynamics (Doebeli 2002). The population size *n*_*i*_ of type *i* is described by

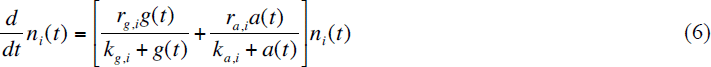

where *r*_*g,i*_ and *r*_*a,i*_ are the maximum growth rates of type *i* on, say, glucose and acetate, respectively; and *k*_*g,i*_ and *k*_*a,i*_ are the resource concentrations that permit growth at half-maximum rate. The concentrations of glucose *g*(*t*) and acetate *a*(*t*) are described by

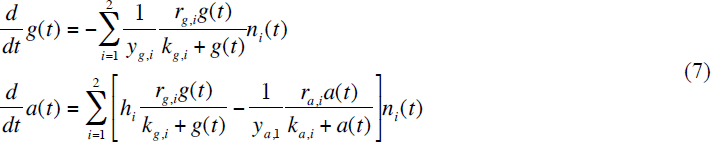

where *y*_*g,i*_, and *y*_*a,i*_ are the population yields from each unit of glucose and acetate, and *h*_*i*_ is the number of units of acetate produced for each unit increment of *n*_*i*_. This system of four equations can be solved numerically to obtain the trajectories of *n*_1_(*t*), *n*_2_(*t*), *g*(*t*), and *a*(*t*) over the course of one or more growth cycles (Fig. 3). This solution depends on 16 parameters, including the initial glucose concentration at each transfer and the dilution factor (which determines the number of generations per growth cycle). Each ecotype may exhibit superior growth on its specialty resource by having a higher maximum growth rate, a lower half-maximum concentration, or some combination thereof. In this picture, the glucose specialist grows faster at the beginning of the growth cycle when the glucose concentration is high, while the acetate specialist grows faster later when the acetate byproduct has accumulated to high concentration.

**Figure 3.**
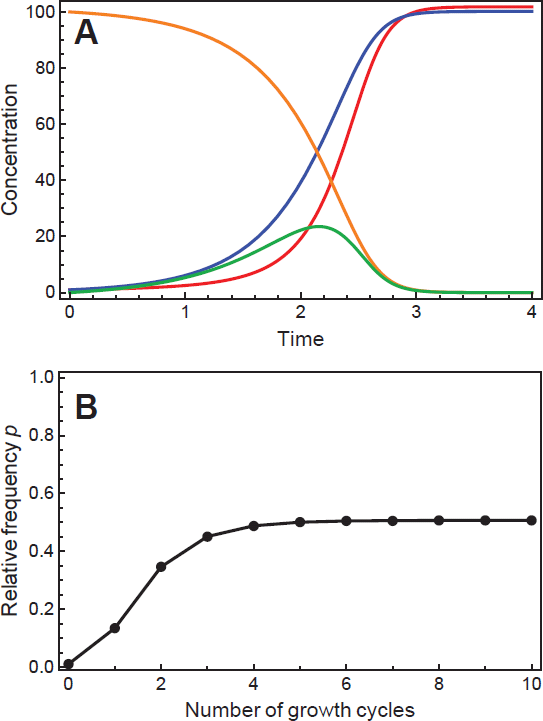
Dynamics of the Monod model of growth and resource concentration for a cross-feeding interaction. (A) Model trajectories of growth and resource concentration over a single growth cycle. Curves show concentrations of an exogenously supplied resource such as glucose (orange), a secondary resource generated as a byproduct such as acetate (green), a glucose specialist ecotype (blue), and an acetate cross-feeding ecotype (red). (B) Relative frequency of the cross-feeding ecotype at the end of each growth cycle during multiple cycles of serial dilution and growth. Parameters used: *r*_*g*,1_ = 1, *r*_*g*,2_ = 2.68, *r*_*a*,1_ = 7, *r*_*a*,2_ = 1, *k*_*g*,1_ = *k*_*g*,2_ = 50, *k*_*a*,1_ = *k*_*a*,2_ = 50, *y*_*g*,1_ = *y*_*g*,2_ = 1, *y*_*a*,1_ = *y*_*a*,2_ = 1, *h*_1_ = *h*_2_ = 1 initial glucose concentration *g*_0_ = 100, dilution factor *F* = 100.

We can use this model to calculate the frequency-dependent relative fitness of two ecotypes in a cross-feeding interaction. At a time point where both resources have been depleted, the model provides a final relative frequency after one growth cycle, which can be analyzed using the same procedure outlined above for quantifying frequency dependence in competition assays. This model neglects mortality during both growth and starvation periods, although model values of *p*_*f*_ are unaffected by mortality provided that each ecotype exhibits the same death rate. As was done with the experimental data in Figs. 1 and 2, we seek values for the unknown parameters *m* and *s* that minimize the mean-squared difference between model values of *p*_*f*_ and the theoretical values of *p*_*f*_ that satisfy Eq. 5. As shown in Fig. S2, solutions to the cross-feeding model are well represented by linear *w*(*p*) for cases of both strong and weak frequency dependence, and with or without the presence of an equilibrium frequency. This indicates that a linear parameterization of *w*(*p*) is the proper null hypothesis for a cross-feeding interaction, and any deviation from linearity would indicate the presence of interactions in addition to those described in Eqs. 6-7.

It is important to note that this resulting function *w*(*p*) does not describe the Monod dynamics of the growth cycle—rather, it describes a population-genetic model (Eq. 3) that results in a post-growth cycle ecotype ratio equal to that produced by the Monod model (Eqs. 6-7). The function *w*(*p*) does not correspond to the Monod dynamics, because the growth rate of each type at any given time is not strictly dependent on its frequency, but rather on the current concentrations of glucose and acetate. Therefore, frequency-dependent fitness in batch culture can only be defined either as an average over an entire growth cycle, or as an “effective” theory that yields results identical to those of an explicit model. In this case, the benefit of an effective theory is that the measured *w*(*p*) can be applied to simple Wright-Fisher type models of allele frequency dynamics.

Although this illustrative example is specific to the cross-feeding interaction, an identical form of frequency dependence is expected under a Black Queen scenario (Morris et al. 2012; Morris et al. 2014), in which one ecotype benefits from the removal of a toxin or growth-inhibitor by another ecotype (Lenski and Hattingh 1986) (Supporting Information; Figs. S3, S4). This equivalence suggests that the linearity of frequency-dependent fitness is broadly applicable to many microbial interactions.

## Discussion

We have presented a detailed procedure for accurately quantifying the relative fitness *w*(*p*) of two ecotypes as an explicit function of relative frequency *p*. This representation provides a convenient framework for quantifying frequency dependence, because our model shows that *w*(*p*) takes a linear form for cross-feeding interactions of any strength or balance (Fig. S2).

This framework, which accounts for changing relative fitness over the course of traditional pairwise competition assays, provides only small gains in accuracy when competition causes only small changes in relative frequency. In the *nuoM* example, frequency dependence was fairly weak (*ms* ≈ 0.03), and so relative frequencies changed only slightly even after 6 cycles of transfer and growth. As a result, the linear parameterization of *w*(*p*) differed only slightly from the 6-day competition mean fitness computed in the traditional way (Fig. 1).

The importance of accounting for changing relative fitness becomes apparent when the competitions result in large changes in relative frequency. This occurs either when frequency dependence is strong, or when competitions are run for long times, or both—as was case for our competitions with the S and L ecotypes. In that example, the effect of frequency dynamics was clear, because the mean fitness values measured after 6 and 12 days of competition were substantially different from the quadratic parameterization of *w*(*p*) that accurately generates those values (Fig. 2).

Furthermore, the use of this framework indicates that the ecological interaction between the S and L ecotypes results in a nonlinear form of *w*(*p*), which would not otherwise have been evident from a traditional fitness measurement. This nonlinearity suggests the existence of at least one interaction in addition to the cross-feeding interaction described by Eq. 6-7, which predicts linear *w*(*p*). Indeed, an additional interaction has previously been reported for this population: during stationary phase, the death rate of L cells is significantly increased in the presence of S cells, providing the S type with a benefit (though not a net advantage) when common (Rozen and Lenski 2000; Rozen et al. 2005; Rozen et al. 2009). This interaction is consistent with the observed decreasing steepness of *w*(*p*) at high frequency of S, which results from that added benefit (Fig. 2). Therefore, a more complete picture of the ecological interactions between S and L ecotypes includes two components: *negative* frequency dependence based on cross-feeding and *positive* frequency dependence resulting from allelopathy. In fact, the allelopathic interaction can only persist in a well-mixed population because of the existence of the cross-feeding interaction, because positive frequency dependence alone accelerates the extinction of the minority type (Chao and Levin 1981; Lenski and Hattingh 1986).

As mentioned above, the measurement procedure described here requires as least as many competitions as there are free parameters in the representation of *w*(*p*). However, an optimal experimental design requires more extensive measurements. First, full coverage of the *w*(*p*) curve requires competitions at many different *p*_0_, ensuring that the widest possible range of *p* is sampled by the series of competitions. Furthermore, it is beneficial to run competitions for a long time to maximize the change in relative frequency, which underscores the importance of using this framework when striving for accurate measurements. However, if competitions are run for too long, one competitor may be driven to near extinction (which increases the error of the measured frequency), or the competition may spend the majority of its time near equilibrium (which swamps the intended measurement of selection). Since the experimenter is unlikely have *a priori* knowledge of the ideal competition time, it is beneficial to measure *p*_*f*_ at multiple time points to ensure a measurement at near maximum precision.

Measurement at multiple time points provides one further significant advantage. When fitness is frequency-dependent, there is a tradeoff in precision when running competitions for a long time: long competitions increase the signal/noise ratio, but the allele frequency dynamics that elucidate the frequency dependence are “averaged out” over the course of the competition. So while long competitions provide high-resolution measurements of the mean fitness between *p*_0_ and *p*_*f*_, they can not resolve the fitness at any particular frequency as well as a shorter measurement could. Measuring *p*_*f*_ at multiple time points alleviates this tradeoff: by fitting a set of parameters that must be consistent with multiple time points, the model is tightly constrained and results in maximum precision. For example, using 6-day and 12-day competitions in combination, we were able to show definitively that a quadratic representation of *w*(*p*) is better than a linear representation (ΔAIC = 15.3, odds ratio = 2130), while this conclusion can not be made based on any single time point (ΔAIC = 3.5, odds ratio = 5.8 for 6-day only; ΔAIC = 0.8, odds ratio = 1.5 for 12-day only). In principle, additional time points in the *nuoM* example could also prove that interaction to be nonlinear, although the time-averaging issue in that case is minimal due to comparatively weak frequency dependence.

In conclusion, we have demonstrated the application of a novel framework for measuring frequency-dependent fitness in microbial populations. Using a Monod model of ecotype dynamics, we have justified a linear parameterization for the frequency-dependent relative fitness for typical cross-feeding interactions. We have further demonstrated the quantification of a complex ecological interaction where linear parameterization fails. We emphasize that accounting for changing relative fitness is important for achieving an accurate measurement of frequency-dependent selection, which to our knowledge has not previously been done in the types of experiments with asexual populations considered here. In addition to quantifying cross-feeding interactions (Doebeli 2002), this technique may be useful to test general theories of evolutionary stable states (Bull and Harcombe 2009; Estrela and Gudelj 2010), and to test hypotheses about the causes of fitness variance within populations (Elena and Lenski 1997). In general, we expect this technique to be widely useful for characterizing evolving microbial ecologies, and it will enable the field to introduce frequency-dependent selection into more general theories of population genetics.

## Acknowledgments

We thank Neerja Hajela for laboratory assistance, and Rohan Maddamsetti, Michael Wiser, and Jeffrey Morris for valuable discussion. This work was supported by an NSF grant (DEB-1019989 to R.E.L.) and the BEACON Center for the Study of Evolution in Action (NSF grant DBI-0939454). The authors declare no conflicts of interest related to the work described herein.

